# Design, construction and optimization of a synthetic RNA polymerase operon in *Escherichia coli*

**DOI:** 10.1101/2021.11.05.467461

**Authors:** Joep Houkes, Lorenzo Olivi, Zacharie Paquet, Nico J. Claassens, John van der Oost

## Abstract

Prokaryotic genes encoding functionally related proteins are often clustered in operons. The compact structure of operons allows for co-transcription of the genes, and for co-translation of the polycistronic messenger RNA to the corresponding proteins. This leads to reduced regulatory complexity and enhanced gene expression efficiency, and as such to an overall metabolic benefit for the protein production process in bacteria and archaea. Interestingly, the genes encoding the subunits of one of the most conserved and ubiquitous protein complexes, the RNA polymerase, are not clustered in a single operon. Rather, its genes are scattered in all known prokaryotic genomes, generally integrated in different ribosomal operons. To analyze the impact of this genetic organization on the fitness of *Escherichia coli*, we constructed a bacterial artificial chromosome harboring the genes encoding the RNA polymerase complex in a single operon. Subsequent deletion of the native chromosomal genes led to a reduced growth on minimal medium. However, by using adaptive laboratory evolution the growth rate was restored to wild-type level. Hence, we show that a highly conserved genetic organization of core genes in a bacterium can be reorganized by a combination of design, construction and optimization, yielding a well-functioning synthetic genetic architecture.

## INTRODUCTION

Operons were first described in the 1960s by Jacob and Monod as “a group of genes regulated by a single operator” (1, 2). At present, operons are generally defined as clusters of genes that are co-transcribed as a single polycistronic mRNA. As first observed in the lactose (*lac*) operon of *E.coli* (1, 2), subsequent experimental analyses of bacterial operons revealed that the clustered genes often encode proteins (or RNAs) with related functions. Indeed, comparative genomics analyses corroborated that operons are generally composed of functionally-related proteins (‘guilt by association’), such as enzymes of a metabolic pathway and subunits of a multi-protein complex (3, 4).

Ever since the discovery of the operon organization, the potential evolutionary forces that drive operon formation have been discussed. Several hypotheses have been suggested to explain how operons could potentially contribute to a selective advantage over individual genes: (i) operons contribute to reduction of genome size and to simplification of gene expression control (5), (ii) operons avoid energy loss through appropriate co-transcription and co-translation of functionally-related genes (6), and (iii) operons improve functional horizontal gene transfer (7–9).

In some metabolic pathways and protein complexes, uneven stoichiometries are required. In these cases it has been demonstrated that differential transcription occurs by using multiple promoters (4, 10), while differential translation of the cistrons within the operons is achieved in multiple ways. The rates of translation initiation can varied by tuning the strength of the Ribosome Binding Sites (RBS) and the mRNA secondary structure around the start codon, as well as by translation elongation, through modulating the codon bias (3, 11–13).

It is interesting to note that, despite being one of the most conserved protein complexes in the three domains of life, the genes coding for the subunits of the prokaryotic DNA-dependent RNA polymerase (RNAP) complex are not clustered in a single operon. At present, not a single prokaryotic genome is known in which the RNAP genes are organized as a single operon. Instead, the genes are spread throughout the genome at different loci. However, in bacteria, in archaea and even in the genomes of chloroplasts in photosynthetic eukaryotes (algae and plants), the RNAP genes are typically co-localized in distinct operons with ribosomal genes (Fig. 1).

**Figure 1.**
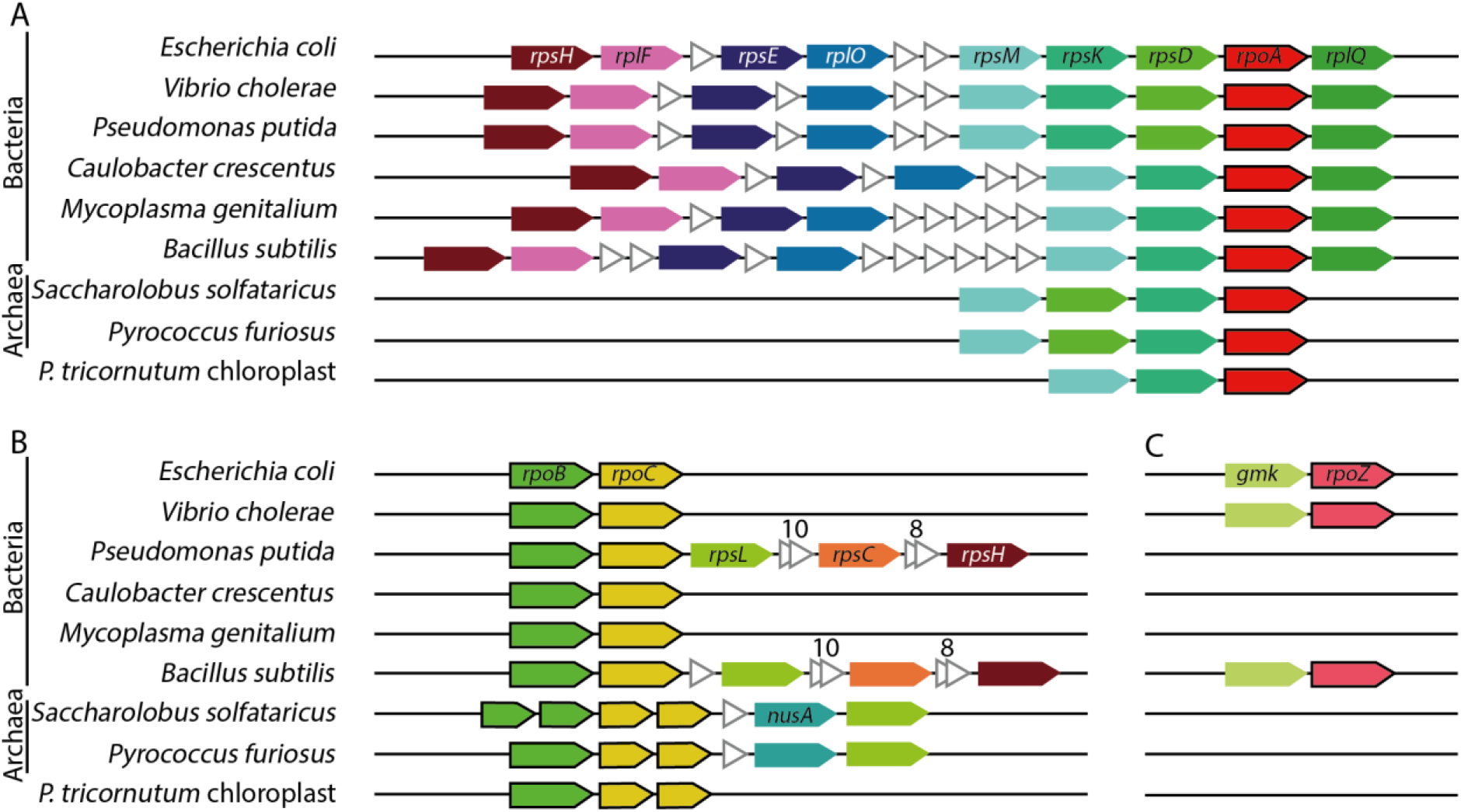
Syntheny of genes encoding core subunits of DNA-dependent RNA polymerase (RNAP). (A) α subunit encoded by *rpoA*, (B) β and β’ subunits encoded by *rpoB* and *rpoC* and (C) ω subunit encoded by *rpoZ* across selected bacterial and archaeal model species, as well as the chloroplast of the microalgae *Phaeodactylum tricornutum*. Same color genes indicate conserved clustering across species. Genes indicated by white triangles are not conserved. *Rps* and *rpl* genes encode ribosomal proteins (16S and 23S subunits, respectively), *nusA* encodes a transcription termination/anti-termination protein and *gmk* encodes a guanylate kinase. Figure generated with data from <u>STRING</u> (string-db.org).

The bacterial RNAP core complex consists of five subunits: two copies of the α subunits and single copies of the β, β’ and ω subunits (14). The *rpoA*-encoded α subunit dimer plays a key role in assembly of the RNAP complex, acting as a scaffold for assembling the β and β’ subunits (15). Furthermore, the α subunit interacts with certain transcription factors to regulate transcription. The *rpoA* gene of *E. coli* and many other bacteria is co-located in an operon harboring ribosomal genes *rpsM, rpsK, rpsD* and *rplQ* (Fig. 1A, (16, 17).

The *rpoB* and *rpoC* genes encode the structurally related β and β’ subunits, respectively, that make up the hetero-dimeric core of the RNAP complex, of which the β’ subunit harbors the catalytic polymerase center (18, 19). Most likely the *rpoB* and *rpoC* genes are the result of a gene duplication (18, 19). In line with this model, a single orthologous RNAP gene still exists in some phages, probably encoding a homo-dimer (19). The bacterial *rpoB* and *rpoC* genes are always clustered, often overlapping, and occasionally fused (20). In addition, a functional synthetic *rpoB-rpoC* fusion protein has been reported (21). In several bacteria the *rpoB* and *rpoC* genes reside in an operon with the ribosomal genes *rplK, rplA, rplJ* and *rplL* (Fig. 1B, 2B). In *E. coli*, this operon has a complex regulation: involvement of four different promoters, regulation by multiple transcription factors and a transcriptional attenuator terminating approximately 70% of transcription just upstream of *rpoB* (Fig. 2B) (22).

**Figure 2.**
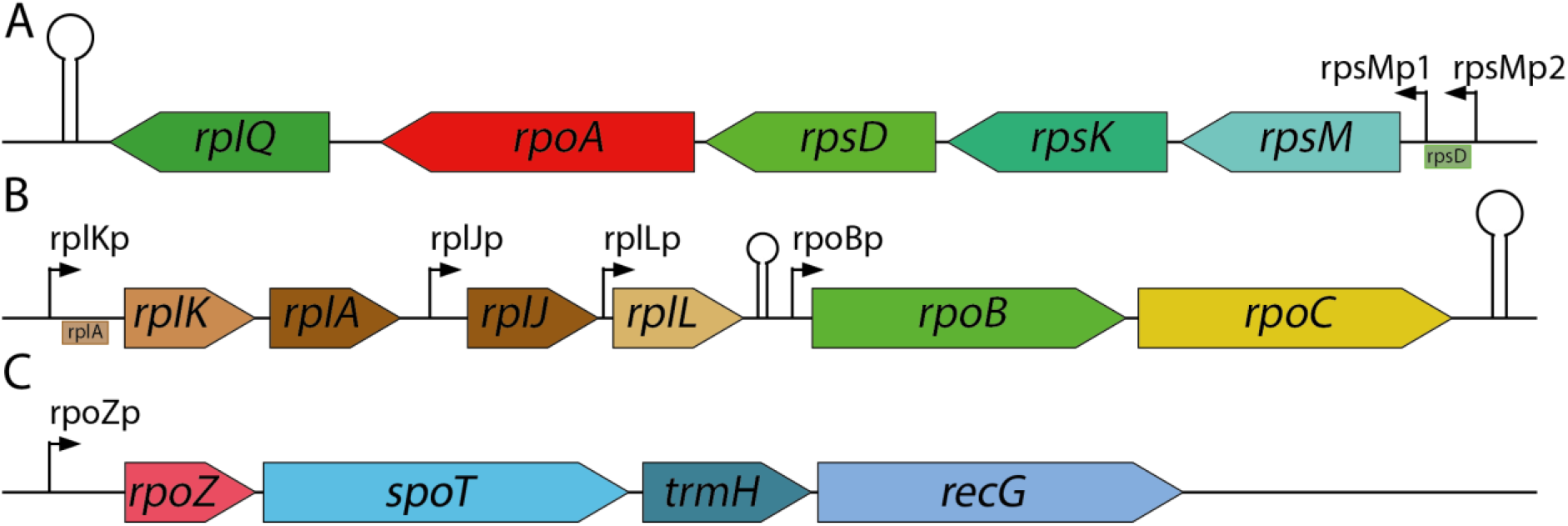
Operons of the RNAP subunits in *E. coli*. (A) operon harbouring the *rpoA* subunit of RNAP, with the ribosomal genes *rpsM, rpsK, rpsD* and *rplQ* (16, 17). (B) operon harboring the *rpoB* and *RpoC* subunits of RNAP, with the ribosomal genes *rplK, rplA, rplJ* and *rplL*. An attenuator between *rplL* and *rpoB* halts approximately 70% of transcription (22). (C) operon harboring the *rpoZ* subunit of RNAP, with *spoT*, involved in stringent stress respons (26–29)*, trmH*, a tRNA methyltransferase (25) and *recG* involved in DNA repair and DNA recombination (24).

The only non-essential subunit of the bacterial RNAP core is the ω subunit, which upon knockout leads to growth retardation, but not to cell death (14, 23). The ω subunit is encoded by *rpoZ*, which in *E. coli* resides in an operon with *trmH, recG* and *spoT* (Fig. 2C). TrmH is a tRNA methyltransferase, and RecG is an ATP-dependent DNA helicase which plays a critical role in DNA repair and DNA recombination (24, 25). SpoT is responsible for the synthesis and degradation of ppGpp, the effector molecule for stringent response, which enables bacterial cells to react to stress conditions by altering expression of many genes (26–29). Interestingly, the primary ppGpp binding site of the *E. coli* RNAP is located at the interface of the β’ and the ω-subunits. The ω subunit plays a role in regulating ppGpp-dependent control of RNAP activity (30), and it has been reported to act as a chaperone for the RNAP subunits (31). The ω subunit binds mainly to the β’ subunit, close to the active polymerase site, indicating a role in controlling the RNAP catalytic activity (18).

The RNAP α_2_ββ’ω core forms a holoenzyme with a σ factor to initiate transcription. Bacteria have several different σ factor, each of which is responsible for transcription of a specific subset of genes (32). The housekeeping σ factor in *E. coli* is σ70, encoded by *rpoD* which controls a large number of promoters, and regulates gene expression during ‘normal’ growth. Six additional σ factors in *E. coli* each control the expression of a particular subset of genes, active during specific environmental conditions (33). Regulation of σ factor expression is very complex, as they are very condition-dependent, unlike the RNAP-core subunits, which are always present.

When comparing the amino acid sequences, the subunit composition, the overall structure, the molecular mechanism and, to some extent, the genomic organization of RNAPs in all domains of life, it becomes apparent they all derive from a common ancestor (34). Both the 13-subunit archaeal and the 12-subunit eukaryotic RNAP complexes contain orthologues of the bacterial RNAP β-, β’-, α- and ω-subunits. This reflects a common evolutionary history, in which the *rpoB/rpoC* gene pair encodes the catalytic β/β’ hetero-dimer of an ancient RNAP variant. At a later stage in the RNAP evolution, the catalytic core was most likely supplemented with the *rpoA*-encoded α-subunit dimer, and the *rpoZ*-encoded regulatory ω-subunit. The archaeal RNAP core resembles the bacterial RNAP complex, with some additional genes encoding auxiliary subunits (34). The three basic eukaryotic RNAPs (Pol I, II, III) and the 2 plant-specific RNAPs (Pol IV, V) are all derived from the archaeal RNAP, each with specific sets of auxiliary subunits (34).

In this study, we set out to use a synthetic biology approach to test if this evolutionary-conserved scattering of the RNAP genes can be reorganized into a single operon, and how such a different architecture would impact cellular fitness. For this, we designed and constructed an operon of the RNAP core genes in *E. coli*. This synthetic operon was introduced on a bacterial artificial chromosome (BAC), and expressed in *E. coli*. Subsequently, native RNA polymerase genes on the *E. coli* chromosome were knocked out, to assess the function of the RNA polymerase operon. This led to a slightly lower growth rate on rich medium compared to wild-type *E. coli*, but, to almost complete loss of growth on minimal medium. However, by adaptive laboratory evolution (ALE) on minimal medium, we could restore growth and even improve the yield of the strain with the synthetic RNAP operon. Overall, this study demonstrates that an evolutionary-conserved operon organization of a core protein complex can be successfully reorganized, suggesting plasticity of genome organization and regulation.

## MATERIAL AND METHODS

### Strains and growth conditions

The *E. coli* DH10B strain (Invitrogen, suppl. table 2) was used for expression of the synthetic operon. This strain was cultured at 37 °C in LB medium (10 g/L tryptone, 10 g/L NaCl, 5 g/L yeast extract), 2xYP medium (16 g/L peptone, 10 g/L yeast extract) or minimal M9+glucose medium (11.28 g/L 5x M9 salts, 0.12 g/L MgSO_4_, 5.5 mg/L CaCl_2_, 3.6 g/l glucose supplemented with 0.1 mL 1000x trace elements solution (50 g/L EDTA, 8.3 g/L FeCl_3_-6H_2_O, 0.84 g/L ZnCl_2_, 0.13 g/L CuCl_2_-2H_2_O, 0.1 g/L CoCl_2_-2H_2_O, 0.1 g/L H_3_BO_3_, 16 mg/L MnCl_2_-4H_2_O) and 0.5mM leucine) at 180 rpm or on LB-agar plates containing 1.5% (w/v) agar (Oxoid) unless stated otherwise. The LB medium was supplemented with different antibiotics (LB/Ab) when appropriate, to final concentrations of 20 μg/mL kanamycin (Carl Roth) (Kan20), or 30 μg/mL chloramphenicol (Sigma Aldrich) (Cam30).

Yeast strain *S. cerevisiae* CEN.PK2-1D (*Euroscarf*, suppl. table 2) strain was used for synthetic operon construction. This strain was cultured at 30°C in 10 mL YPD medium (20 g/L peptone, 10 g/L yeast extract, 20 g/L glucose) or in SC medium (1.9 g/L nitrogen base without amino acids, 5 g/L ammonium sulphate, 20 g/L glucose, 2g/L drop-out mix, appropriate auxotrophic marker (76 mg/L uracil, 380 mg/L leucine, 76 mg/L histidine, 76 mg/L tryptophan)) in 50 mL tubes at 180 rpm.

### Preparation of electrocompetent cells

*E. coli* cells were made electrocompetent by culturing at 37 °C in 2xYP (supplemented with 0.01 M L-arabinose for recombination experiments), typically in 50 mL medium in 250 mL Erlenmeyer flasks, at 200 rpm until an OD600 nm of 0.4 was reached. The cells were then cooled rapidly on ice, and subsequently washed once with 1 culture volume of ice-cold ddH2O and twice with 0.5 culture volumes of ice-cold 10% glycerol. Finally, the cells were suspended in ice-cold 10% (vol/vol) glycerol to a final volume of 200 μL for each 50 mL of initial culture volume.

Electroporation of 20 μL competent cells was performed in ice-cold 2 mm electroporation cuvettes at 2500 V, 200 Ω and 25 μF (ECM 630 BTX). Immediately after electroporation, cell recovery was performed in 1 mL LB medium at 37 °C, 750 rpm for 1 h when plasmids were transformed. During recombination experiments, recovery was performed at 30 °C, 750 rpm for 2.5 h. After recovery, the cells were plated on LB/Ab agar plates. Single colonies were picked and re-suspended in 50 μL of ddH_2_O for colony PCR and used to inoculate 10 mL of LB/Ab for overnight incubation and subsequent isolation of plasmids, using GeneJET Plasmid MiniPrep Kit (Thermo Fisher Scientific). The provided protocol was adjusted by initially centrifuging the cell cultures at 4700 rpm for 10 min, and introducing of an incubation (2 min) at room temperature after addition of Elution Buffer or ddH_2_O (warmed to 70 °C).

Transformations of *S. cerevisiae* were performed by chemical transformation. Cells were plated from glycerol stock on YPD, a single colony was picked for overnight culturing in YPD (typically in 10 mL medium in 50 mL tubes, at 180 rpm). The culture was diluted to OD600nm 0.4 in YPD and incubated at 30°C, 200 rpm for 3 hours (typically in 50 mL medium in 250 mL Erlenmeyer flasks). The cells were then washed with 0.5 culture volume of ddH_2_O. Cells were resuspended in ddH_2_O to a final volume of 1 mL, aliquoted into 100 μL volumes and stored at 4°C for up to a week. To a 100 μL cell suspension, 350 μL of a transformation mix (consisting of 240 μL PEG-3350, 36 μL 1 M LiOAc, 50 μL 2 mg/mL denatured salmon sperm DNA, and 34 μL of DNA mix containing 500 ng of DNA fragments and 1 μg of backbone plasmid) was added. Next, the cells/transformation mix was heat-shocked at 42 °C for 40 min. The cells were resuspended in 1 mL YPD and 500 μL was used to inoculate 5 mL YPD for overnight recovery, to boost the recombination efficiency. The remaining 500 μL was plated on SC-agar plates with the appropriate auxotrophy markers. The success of the transformation was assessed by colony PCR and single colonies were used to inoculate 10 mL SC with the appropriate auxotrophy markers. For plasmid extraction, the cells were resuspended in 200 μL GeneJET Plasmid MiniPrep (ThermoFisher Scientific) resuspension buffer supplemented with 3 μL 1000 U/mL Zymolase (Amsbio) and incubated for 30 min at 37 °C to digest the cell walls. The rest of the extraction was performed according to the MiniPrep protocol.

### Plasmid construction

All PCR reactions for cloning purposes were performed using Q5® High Fidelity 2X Master Mix (New England Biolabs). The reactions mixtures were prepared using 1 ng of template, 25 μL Q5® High Fidelity 2X Master Mix, primers to a final concentration of 500 nM, and ddH_2_O to a final volume of 50 μL (primers are listed in Suppl. table 4.). Amplification products were run on 0.7% agarose gels stained with SYBR Safe DNA Gel Stain (Invitrogen). The bands of interest were then excided, and the DNA purified using Zymoclean Gel Recovery kit (Zymo Research). Cloning was done using HiFi assembly (New England Biolabs) according to manufacturer protocol. Overhangs for HiFi assembly were added as 5’ extensions of PCR-primers (list of plasmids in Suppl. table 3, list of primers in Suppl. table 4).

### Preparation of knock-out strains

Cells were transformed with a linear knock-out fragment with 50 bp recombination flanks harboring a chloramphenicol resistance marker flanked by mutant *lox66* and *lox72* sites that upon recombination are not recognized anymore by Cre recombinase (35). After recombination, single colonies were picked, streaked on LB/Ab agar plates supplemented with isopropyl-β-D-thiogalactopyranoside (IPTG) (Fisher Scientific, catalogue) to a final concentration of 0.5 mM, and then re-suspended in ddH_2_O to perform colony PCR to determine whether appropriate recombination occurred. IPTG-induced expression of Cre recombinase from the BAC vector, generating mixed colonies with chloramphenicol resistant (CmR) and sensitive (CmS) cells. These mixed colonies were re-streaked on LB/Ab plates. Single colonies were picked from these plates and re-streaked on LB agar and in LB agar/Cam30. CmS colonies were picked and re-suspended in ddH_2_O to perform colony PCR, to confirm the successful excision of the chloramphenicol resistance gene.

### Growth assays of knock-out strain and data analysis

After knock-out was confirmed by colony PCR, growth assays were performed to assess growth rates of the mutant strains. Precultures were prepared on LB for each strain and for wild-type DH10B. Precultures were washed 3 times with ddH_2_O and diluted to OD600 0.1. Next, 15 μL of diluted preculture of each strain and 135 μL of LB or M9+Glucose was transferred in a 96-well plate. The wells were covered with 50 μL of mineral oil (Bio-Rad), to avoid evaporation during the experiment. The plate was then incubated at 37 °C in a Biotek ELx800 absorbance microplate reader (Fisher Scientific). The provided reader control software, Gen5, was used to set a measuring protocol consisting of a cycle of 5 min of linear shaking followed by absorbance measurement at 600 nm, for at least 24 h. The data were then exported to an Excel spreadsheet. An in-house MatLab script was used to process the data, yielding strain-specific growth graphs and doubling times.

### sequencing and analysis

For genomic sequencing analysis DNA was isolated and sequenced using Illumina NovaSeq paired end 150bp. Mutation analysis was done using BreSeq(36) using the DH10B genome (NCBI ref.: NC_010473.1) as reference. In addition, Genious Prime was used to map the sequencing results to a DH10B reference genome and to validate the genomic knockouts of the four RNAP genes.

## RESULTS AND DISCUSSION

### Designing and building a synthetic RNAP operon

We rationally designed a synthetic RNAP operon in the order *rpoABCZ*. First, *rpoA* was introduced by Gibson assembly on bacterial artificial chromosomes (BACs), controlled by a few different promoters and RBS. One variant contained the native *rpoA* promoter, i.e. the rpsMp2 promoter of the operon, and the native *rpoA* RBS. In addition, *rpoA* was inserted downstream three constitutive promoters of different strength (weak, moderate, strong) (37), that each were combined with one of 5 RBS in a linearly increasing strength range (20, 40, 60, 80 and 98% of the predicted maximum strength), as designed by EMOPEC (38). For each combination, the native *rpoA* gene on the *E. coli* chromosome was knocked out, and a comparative growth assay on LB medium determined the best performing combination: strong constitutive promoter and RBS80 (Suppl. table 1). Interestingly, knock-out of the native *rpoA* gene was successful only for 7 out of 24 promoter-RBS combinations, strongly suggesting that there is a certain range of *rpoA* expression levels that allows for *E. coli* survival. To allow for easy addition and selection of subsequent subunits, we decided to move the marker directly downstream of the operon. The *kanR* resistance marker was removed from the BAC-*rpoA* using the λ-red system and *cre* recombinase. This created an addictive plasmid bale to be propagated because of the presence of the essential *rpoA* gene. We then introduced *rpoB* with 6 different RBSs (native and 5 synthetic variants) and a *kanR* gene directly downstream of *rpoA*. Deletion of chromosomal *rpoB* gene and subsequent growth assays of the 6 RBS variants, showed that the native RBS associated with *rpoB* on the BAC, resulted in the fastest growth. We aimed to continue this approach for *rpoC* as well, but several attempts to introduce *rpoC* in the operon on the BAC were not successful. To introduce *rpoC* we aimed to switch the antibiotic resistance gene in the BAC to *tetR*, to select for BAC-*rpoABC* after transformation into the double knockout strain. Unexpectedly we did not obtain any transformants harboring *rpoABC*. This prevented us from following the planned approach, to properly introduce, delete and optimize expression for each RNAP gene one-by-one.

Therefore, an alternative approach was designed. As we could not continue with the one-by-one optimization of the *rpo* genes, we decided to assemble the operon at once. Making a combinatorial library of the synthetic *rpo* operon would lead to a large collection of *E. coli* strains from which all genomic *rpo* genes would first need to be deleted and confirmed, leading to a major experimental effort. Hence, we decided to use the previously identified well performing strong promoter in combination with RBS80 for *rpoA*, and the native RBS variant for *rpoB*. In addition, without prior knowledge, we tested the RBS80 variant upstream *rpoC* as well as upstream *rpoZ*. Downstream rpoZ, we included an in-house designed synthetic terminator consisting of a <u>stem</u>-loop and a T-stretch (5’-<u>ccccgc</u>ttcg<u>gcgggg</u>ttttttt) (Fig. 3). To efficiently assemble all these parts in the relatively large BAC construct at once, we chose to further construct the BAC in *Saccharomyces cerevisiae* because of its highly efficient recombination system. For that purpose, a BAC-yeast artificial chromosome (BAC-YAC) shuttle vector was constructed. First, we PCR amplified the bacterial replication system (*sopA, sopB, sopC* and *repE*) from a BAC (pBeloBAC11 (39)), as well as the yeast centromere region from a YAC (pHLUM (40)) with a *his3* and a *ura3* selection marker. Furthermore, the RNAP genes *rpoA, rpoB, rpoC* and *rpoZ* (and the aforementioned RBS variants) were PCR-amplified from *E. coli* DH10B. To allow for eventually knocking out the native *rpo* genes, the genes encoding the λ-red system (*gam, bet, exo*) and the Cre recombinase (*cre*) were PCR amplified by using plasmid pSC020 (41) as a template. All PCR amplifications were carried out using extended primers that generated specific 50 base pair overhangs to allow for efficient homologous recombination in *S. cerevisiae*. Next, we transformed *S. cerevisiae* CEN.PK2-1D (histidine, leucine, tryptophan, uracil auxotroph) with the 8 fragments as described above for homologous recombination and selected for correct assemblies using medium lacking histidine and uracil (Fig. 3). The resulting colonies were demonstrated by PCR to contain the designed BAC-YAC clone. Two of these colonies were used for plasmid isolation and transformation into *E. coli* DH10B. From two *E. coli* DH10B transformants, the sequence of the obtained BAC-YAC constructs was analyzed and confirmed to be correct.

**Figure 3.**
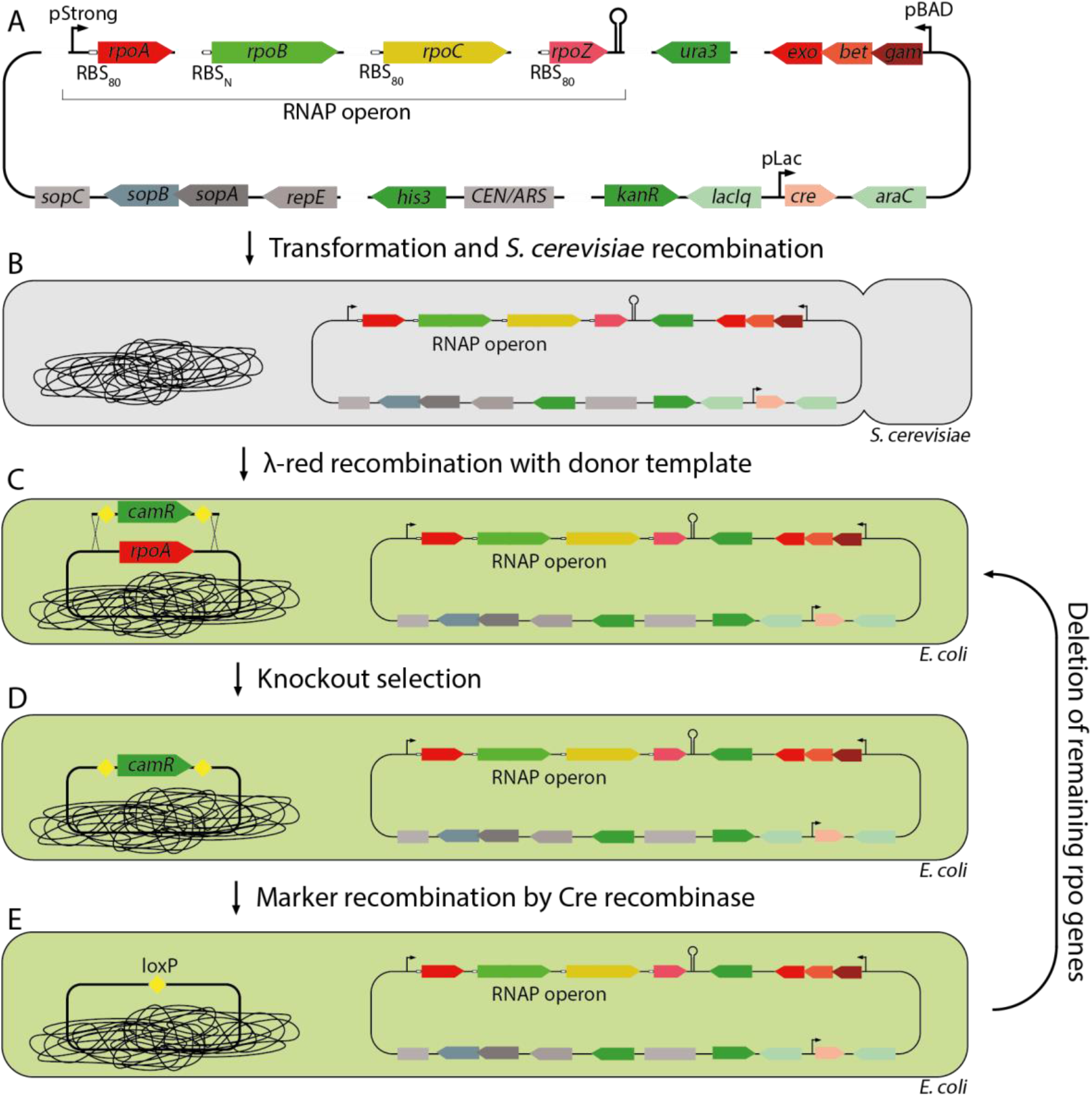
Design and construction of RNAP operon on BAC-YAC and knockout of native genes. (A) Design of BAC and parts utilized for assembly. (B) Following this, yeast recombination, isolation and transformation into *E. coli* DH10B. (C) Knockouts are performed using λ-red recombination, (D) after which selection occurs using the antibiotic resistance marker. (E) Finally, the antibiotic resistance marker is recombined using Cre recombinase, leaving a dysfunctional *loxP* scar. Steps C-E are repeated for remaining *rpo* genes.

### Knockout strategy

After transformation of *E. coli* DH10B with the BAC-YAC shuttle vector harboring the RNAP operon, the native genes were knocked out using λ-red recombination (Fig. 3). For this, a repair template containing a chloramphenicol resistance gene flanked by *lox66* and *lox72* sites was used (35). The repair template was PCR-amplified using primers harboring overhangs homologous to the knockout location. Using this approach, the chromosomal *rpoA, rpoB-rpoC* and *rpoZ* genes of the BAC-YAC-containing *E. coli* strain were deleted consecutively (Fig. 3). The successful genomic deletions were confirmed initially by PCR, and finally by genome sequencing. This confirmed that the synthetic operon could fully replace the scattered genomic *rpo* genes.

To assess the growth of this newly created strain, named strain JH10B, growth assays were performed on rich medium (LB) and minimal medium (M9+glucose),respectively (suppl. fig. 1). It was found that on rich medium the growth rate of JH10B was slightly lower (reduced growth rate approximately 7%) compared to growth of the wild-type *E. coli* DH10B. On minimal medium, however, no growth was observed within the first two days for the engineered strain harboring the RNAP operon (suppl. Fig. 1).

### Evolutionary optimization

In an attempt to recover the ability of strain JH10B to grow on minimal medium, we decided to perform adaptive laboratory evolution (ALE) on M9+glucose. For this, two colonies (biological replicates a & b) were randomly selected and grown in 10 mL M9+glucose until the OD600 was at least 0.4. After this, 10 μL (0.1%) was transferred to a fresh tube with 10 mL M9+glucose, and after each passage a sample was taken for storage. After the first passage the obtained strain JH10B-ALE-1 already started growing on M9+glucose, and after 12 passages (strain JH10B-ALE-12) a plate reader experiment was done to assess growth of selected generations of the adapted strains (JH10B-ALE-1,-2,-3,-4,-8,-12) in minimal and rich medium, compared to the wild-type strain (Fig. 4, suppl. fig. 2). Although the lag-phase of the wild-type is shorter than the lag phases of the ALE strains, the growth rate of the evolved strains is higher (up to 23%). Already after one round of ALE, the doubling time of both biological ALE-1 replicates (strains JH10B-ALE-1a/b; 2.2/1.8 hrs respectively) was comparable to the wild-type strain (1.6 hrs), whereas after the second round of ALE the evolved strain (JH10B-ALE-2b) grows faster than the wild-type on M9+glucose medium (1.4 hrs). Additionally, the yield (final OD600) of some of the evolved strains (JH10B-ALE3b; 1.39) are substantially higher compared to the yield of the wild-type strain (0.66) (suppl. fig. 2). While this experiment enhanced the growth rate of JH10B on M9+glucose, growth on LB did decrease (approximately 42% for JH10B-ALEa and 35% for JH10B-ALEb) during the experiment (Suppl. fig. 3).

**Figure 4.**
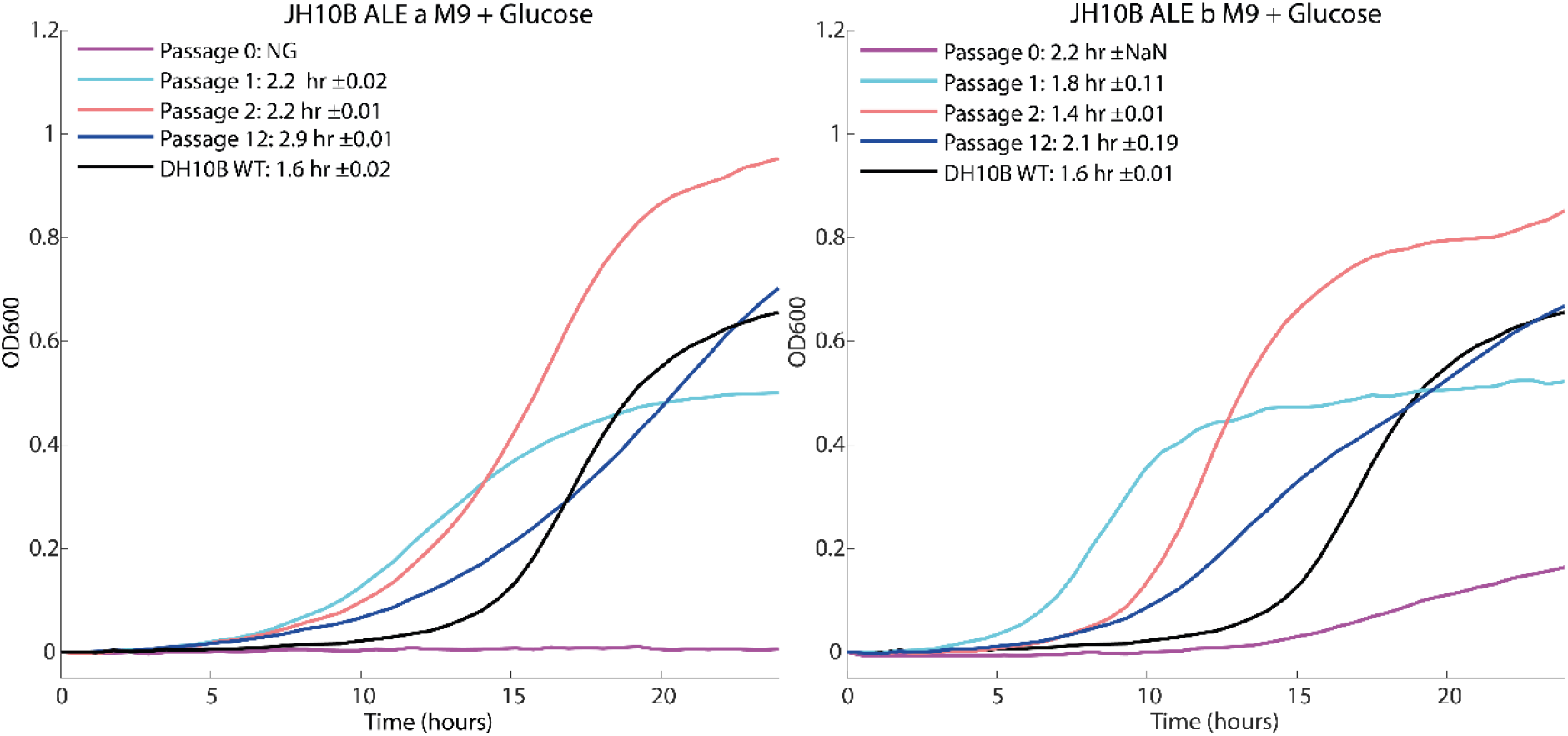
**Growth assays of *E. coli* strains harboring the RNAP operon after ALE** on M9 medium with glucose. Representation of each line shown in figure legend. Doubling time and standard deviation of 6 replicates are indicated.

### Genome sequencing and mutational analysis

Whole-genome sequencing was performed on both biological replicates of the strains obtained after one adaptation cycle (JH10B-ALE1a and JH10B-ALE1b), with the wild-type (passage 0) as control. In total one unique mutation was found in *rpoB* in JH10B-ALE1a (T563P) and one unique mutation in *rpoC* in JH10B-ALE1b (128LDMPL duplication), compared to the parental strain. Another mutation in *rpoA* (Y68C) was found not only in both ALE1 replicates, but also in passage 0, before ALE (Table 1). Sequencing results indicate that this mutation appeared either during PCR amplification or during yeast recombination/replication of the fragment. Visual inspection of the crystal structure indicates that the duplication in *rpoC* is located at the interface of the DNA strand. The mutation in *rpoA* in both replicates localizes in a loop close to the binding site with the β and β’ subunits.

**Table 1.**
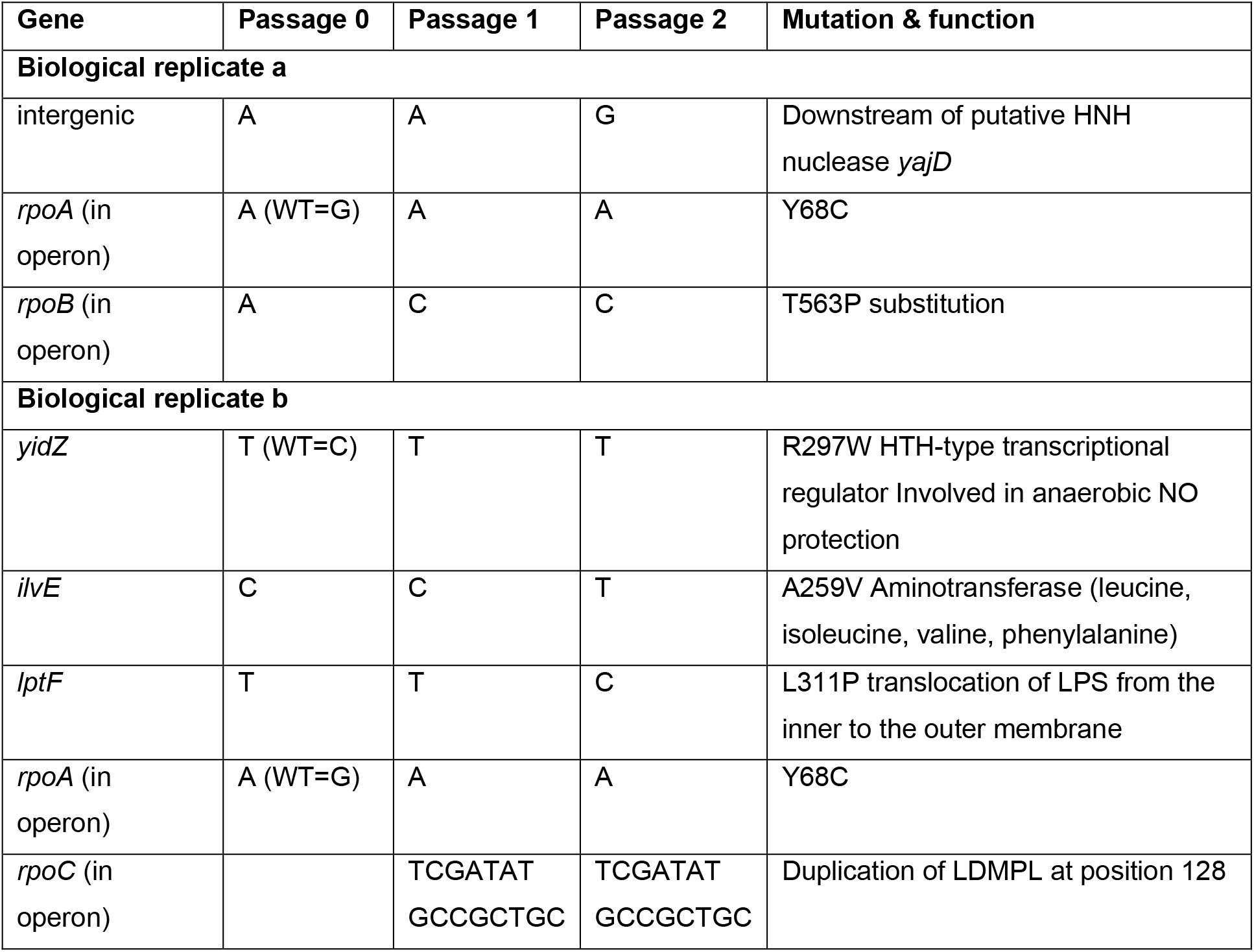
mutations in *E. coli* strains harbouring RNAP operon after ALE experiment.

Interestingly, but perhaps not surprisingly, during previously performed ALE experiments with *E. coli*, mutations are frequently found in the genes encoding the RNAP subunits (42–47). In one of these studies it has been concluded that mutations in the RNAP complex can generally satisfy selection of enhanced growth rates under many conditions (47). Being the most central transcriptional regulatory hub, many amino acid substitutions at relevant sites in the RNAP complex can potentially lead to differences in the host’s transcriptional profile (48, 49). In one study, 80% of *E. coli* MG1655 strains that were adapted to a minimal medium with glycerol as carbon source, appeared to harbor mutations in the *rpoC* gene (46). These *rpoC* mutations appeared to lead to a 60% increase in growth rate in glycerol minimal medium after reintroduction to MG1655 while simultaneously changing the expression pattern of 900-1200 (20–27%) of its genes. At the time of writing, in ALEdb (www.ALEdb.org), a web-based platform that reports on published ALE-acquired mutations for *E. coli* contains 21738 unique mutations. Of these, 132 were found in *rpoB*, 109 in *rpoC* and 47 in *rpoA* and none in rpoZ, for a total of 288 unique mutations, 1.32% of all unique mutations from the database, while these genes make up 0.21% of the *e coli* genome(50). The mutation found in *rpoB* in this study (T563P) has been found before during ALE on minimal medium, this mutation could confer rifampicin resistance(47).

Although selection for enhanced growth under specific conditions very often leads to mutations in RNAP genes, this often comes with a cost. While the fitness increases in the selected environments, it decreases in different environments (46, 51, 52). In the present study, ALE on M9 glucose led to faster growth in this condition (Fig. 4), but to reduced growth on LB medium (suppl. fig. 3). The obtained amino acid substitutions can have different effects on the subunit, and on the RNAP complex. Some studied mutations in *rpoC* have been shown to decrease open complex half-life (46, 53), which affects the transcription initiation and elongation activity (54). These studies also show that down-regulated genes often have promoters with stress-related σ factors, whereas up-regulated genes rather tend to have growth-related σ factors (53). It is not exactly known how mutations in RNAP genes modulate the expression of genes that lead to certain phenotypes. However, the RNAP complex can be regarded as the ultimate transcriptional regulator, allocating the cellular resources to the specific molecular functions.

### Lessons for synthetic genome re-organization and modularization

Based on the here presented successful transplantation of separate chromosomal genes to a fully functional BAC-based RNAP operon, we conclude that, at least under the tested conditions, the ubiquitous ‘coupling’ of the bacterial and archaeal RNA polymerase genes with the ribosomal genes is not essential for life. This is an encouraging result for the design and construction of synthetic genomes, based on partial rational combinations of components. Some attempts to construct rearranged synthetic genomes have already been made, most notably in a minimized *Mycoplasma* species JCVI-Syn3.0(55) and in *S. cerevisiae* (56). In *S. cerevisiae*, a core set of 13 glycolysis genes, which in nature are scattered across the yeast chromosomes, were expressed from one chromosomal locus using the native promoters, after which the native genes were deleted. This led to a strain which was phenotypically similar to the wild-type strain. This demonstrated that co-localization of genes is feasible even for a eukaryotic species with multiple chromosomes. In the prokaryotic minimal cell JCVI-Syn3.0, genome reorganization towards clustering of functionally related genes was attempted as well. First, genes were split in seven different categories for co-localization (e.g. DNA repair, transcription, translation, glycolysis). Then the genome was split in eight segments, and for each segment separately all genes belonging to one category were co-localized within this segment. However, this meant that some original operon structures had to be rearranged. Hereto, a relatively random approach was followed by which promoters/RBS were manually selected without any optimization to regulate the reorganized genes. This highly randomized approach still led, maybe surprisingly, to viable cells after modularization of one out of eight segments. However, the other seven reorganized segments did not lead to viable cells. The latter result strongly suggests that a rational approach should probably be combined with a random/combinatorial approach as described in this study, to allow for selecting appropriate combinations of promoter and RBSs.

In conclusion, the current study reveals how modularization in bacteria can be performed via a step-wise introduction of genes with synthetic control elements (promoters, RBSs) and subsequent deletion of the native genes. This approach could be the basis to modularize larger parts of the genome. After introduction of a synthetic gene or operon with a small library of synthetic promoters or RBS, the native gene(s) can be deleted to identify viable clones, which could optionally be further improved by ALE. This strategy will take relatively long, but could be speeded up lab-automation. Eventually this may lead to a fully reorganized modular genome, that could be highly beneficial for easy ‘swapping’ of modules when engineering cells towards desired applications.

## AVAILABILITY

All data included in this study is available upon request by contact with the corresponding author.

## ACKNOWLEDGEMENT

We thank dr. Sjoerd Creutzburg for supplying plasmid pSC020 and the synthetic terminator, and dr. Arren Bar-Even for supplying the MatLab script for determining growth rates.

