## Supplementary Data for "Design, construction and optimization of a synthetic RNA polymerase operon in *Escherichia coli*"

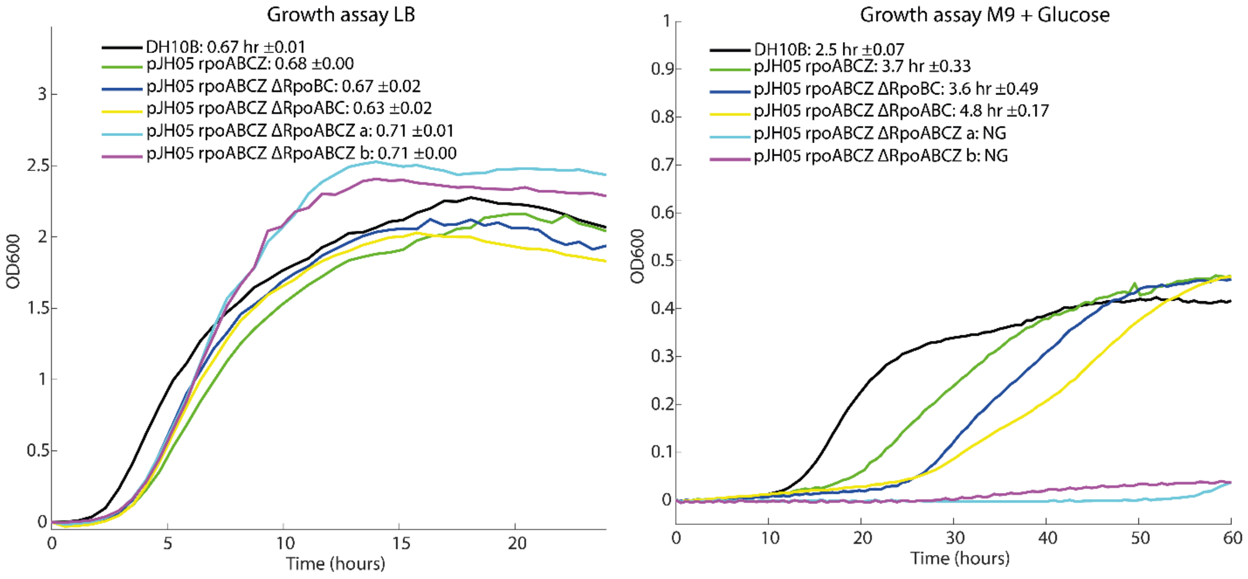


**Suppl. Fig. 1. Growth assays of *E. coli* strains harboring the RNAP operon** on LB (left) and M9 medium with glucose (right). Representation of each line shown in figure legend, numbers are doubling time, time point of highest doubling time, and max OD reached, respectively. Lines are means of 6 technical replicates.

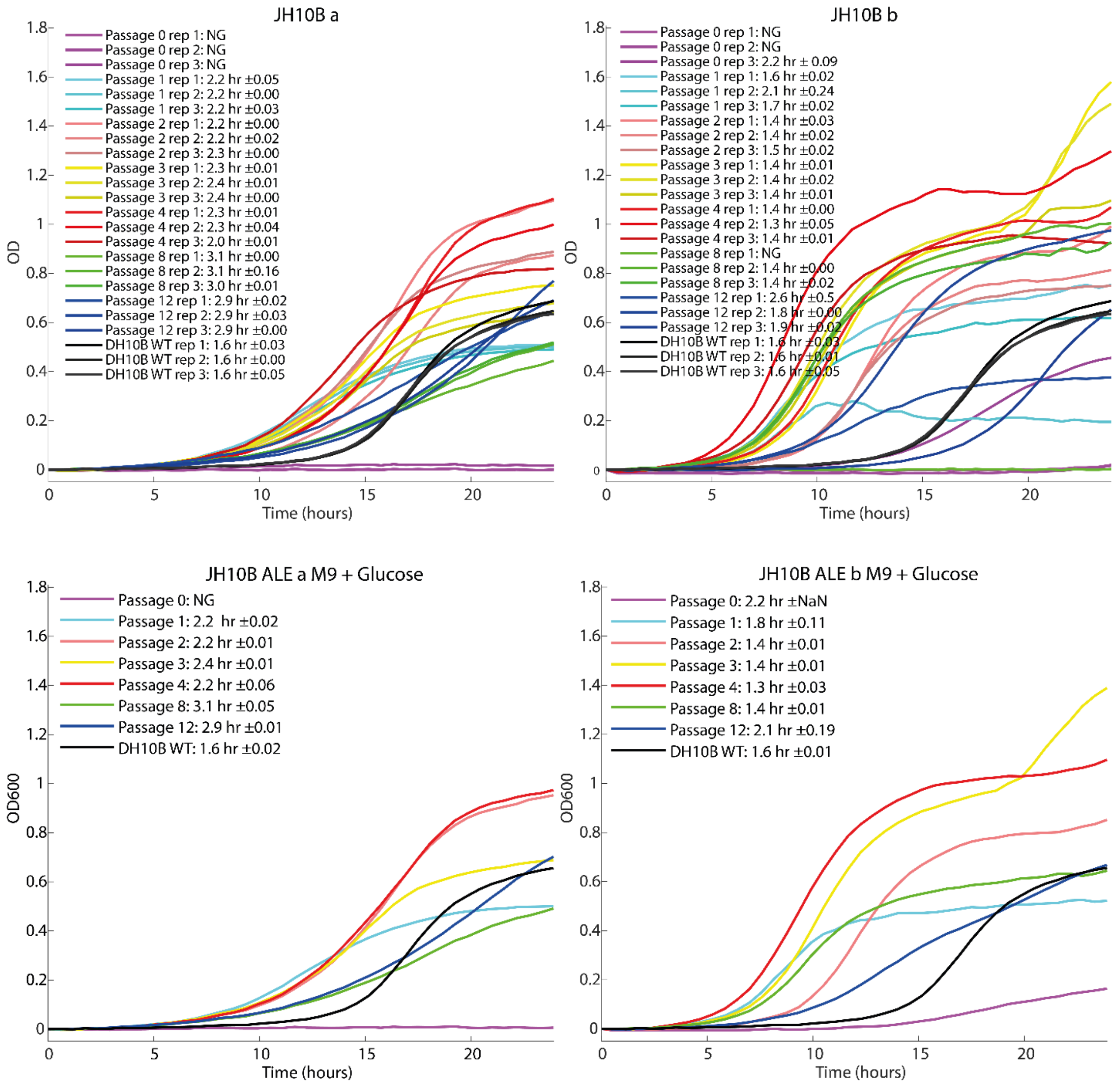


**Suppl. fig. 2. Growth assays of E. coli strains harboring the RNAP operon after ALE on M9+glucose, all replicates**. Top graphs has all three biological replicates per passage consisting out of 2 technical replicates split out. Bottom graphs have biological replicates combined. Each Representation of each line shown in figure legend. Doubling time and standard deviation are indicated.

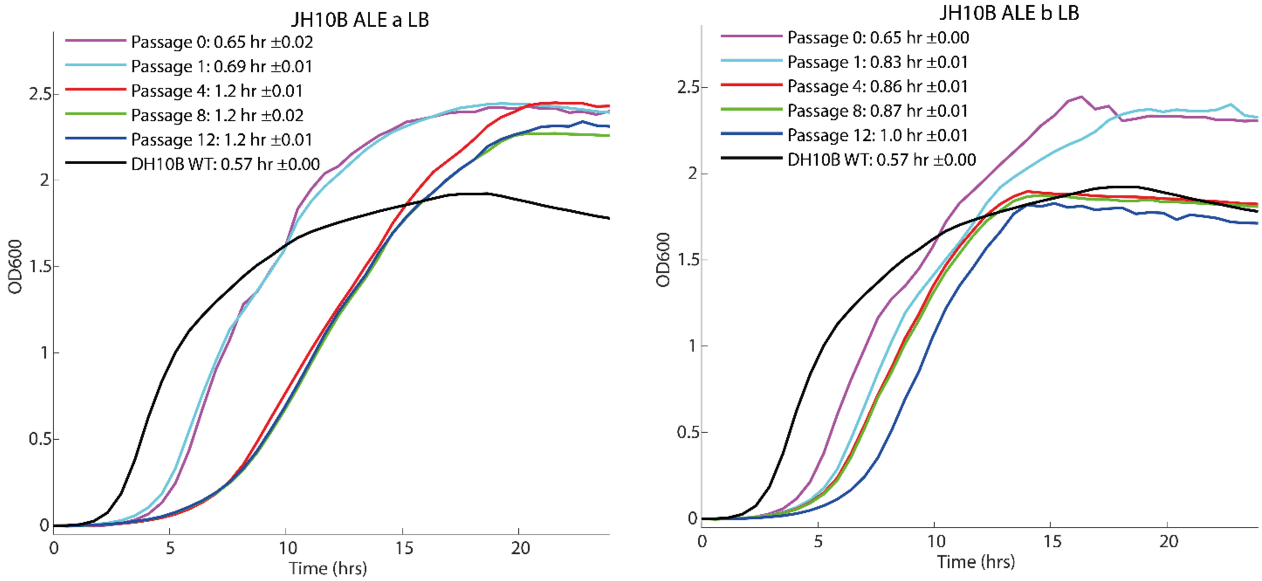


**Suppl. Fig. 3 Growth assays of E. coli strains harboring the RNAP operon after ALE on LB medium**. Representation of each line shown in figure legend. Doubling time and standard deviation of 6 replicates are indicated.

**Suppl. table 1. Combinations of promoters and RBSs tested for *rpoA*.** Combination in red was not obtained, green combinations allowed complementation of the lethal Δ*rpoA* mutation, doubling times measured in the consequent growth assay are included.

| | $P_{Weak}$ | $P_{Moderate}$ | | $P_{Strong}$ | | $P_{Native}$ | |
|---|---|---|---|---|---|---|---|
| $RBS_{20}$ | pJL_wA20 | pJL_mA20 | | pJL_sA20 | | pJL_nA20 | |
| $RBS_{40}$ | pJL_wA40 | pJL_mA40 | | pJL_sA40 | | pJL_nA40 | |
| $RBS_{60}$ | pJL_wA60 | pJL_mA60 | | pJL_sA60 | 0.54 | pJL_nA60 | |
| $RBS_{80}$ | pJL_wA80 | pJL_mA80 | 0.553 | pJL_sA80 | 0.539 | pJL_nA80 | |
| $RBS_{98}$ | pJL_wA98 | pJL_mA98 | | pJL_sA98 | | pJL_nA98 | |
| $RBS_{N}$ | pJL_wAN | pJL_mAN | 0.549 | pJL_sAN | 0.544 | pJL_nAN | 0.543 |

**Suppl. table 2. List of strains**

| Strain | Comment | Source |
|---|---|---|
| *E. coli* DH10B | Used for cloning purposes | Invitrogen |
| *E. coli* DH10B pJL_mA80 Δ*rpoA* | DH10B Δ*rpoA* harboring pJL_mA80 | This work |
| *E. coli* DH10B pJL_mAN Δ*rpoA* | DH10B Δ*rpoA* harboring pJL_mAN | This work |
| *E. coli* DH10B pJL_sA60 Δ*rpoA* | DH10B Δ*rpoA* harboring pJL_sA60 | This work |
| *E. coli* DH10B pJL_sA80 Δ*rpoA* | DH10B Δ*rpoA* harboring pJL_sA80 | This work |
| *E. coli* DH10B pJL_sAN Δ*rpoA* | DH10B Δ*rpoA* harboring pJL_sAN | This work |
| *E. coli* DH10B pJL_nAN Δ*rpoA* | DH10B Δ*rpoA* harboring pJL_nAN | This work |
| *E. coli* DH10B pJH05 Δ*rpoA* | DH10B Δ*rpoA* harboring pJH05 | This work |
| *E. coli* DH10B pJH05 Δ*rpoA* Δ*rpoB/rpoC* | DH10B Δ*rpoA* Δ*rpoB/rpoC* harboring pJH05 | This work |
| *E. coli* JH10B | DH10B Δ*rpoA* Δ*rpoB/rpoC* Δ*rpoZ* harboring pJH05 | This work |
| *E. coli* JH10B 1-12 a | DH10B Δ*rpoA* Δ*rpoB/rpoC* Δ*rpoZ* harboring pJH05. Evolved for 1-12 generations in M9+glucose | This work |
| *E. coli* JH10B 1-12 b | DH10B Δ*rpoA* Δ*rpoB/rpoC* Δ*rpoZ* harboring pJH05. Evolved for 1-12 generations in M9+glucose | This work |
| *S. cerevisiae* CEN.PK2-1D | MATa/α ura3-52/ura3-52 trp1-289/trp1-289 leu2-3_112/leu2-3_112 his3 Δ1/his3 Δ1 MAL2-8C/MAL2-8C SUC2/SUC2 | Euroscarf |

**Suppl. table 3. List of plasmids**

| Plasmid | Description and relevant characteristics | Reference |
|---|---|---|
| pHLUM | Harbors CEN/ARS, *his3* and *ura3* used in shuttle vectors | (1) |
| pBeloBAC11 | Harbors BAC replication system *sopA, sopB, sopC* and *repE* | (2) |
| pSC020 | Harbors λ-red system and *cre* | (3) |
| pJL_wA20 | BeloBAC backbone, λ-red system, *cre*, *kan* and *rpoA* under weak promoter and $RBS_{20}$ | This work |
| pJL_wA40 | BeloBAC backbone, λ-red system, *cre*, *kan* and *rpoA* under weak promoter and $RBS_{40}$ | This work |
| pJL_wA60 | BeloBAC backbone, λ-red system, *cre*, *kan* and *rpoA* under weak promoter and $RBS_{60}$ | This work |
| pJL_wA98 | BeloBAC backbone, λ-red system, *cre*, *kan* and *rpoA* under weak promoter and $RBS_{98}$ | This work |
| pJL_wAN | BeloBAC backbone, λ-red system, *cre*, *kan* and *rpoA* under weak promoter and $RBS_{N}$ | This work |
| pJL_mA20 | BeloBAC backbone, λ-red system, *cre*, *kan* and *rpoA* under medium promoter and $RBS_{20}$ | This work |
| pJL_mA40 | BeloBAC backbone, λ-red system, *cre*, *kan* and *rpoA* under medium promoter and $RBS_{40}$ | This work |
| pJL_mA60 | BeloBAC backbone, λ-red system, *cre*, *kan* and *rpoA* under medium promoter and $RBS_{60}$ | This work |
| pJL_mA80 | BeloBAC backbone, λ-red system, *cre*, *kan* and *rpoA* under medium promoter and $RBS_{80}$ | This work |
| pJL_mA98 | BeloBAC backbone, λ-red system, *cre*, *kan* and *rpoA* under medium promoter and $RBS_{98}$ | This work |
| pJL_mAN | BeloBAC backbone, λ-red system, *cre*, *kan* and *rpoA* under medium promoter and $RBS_{N}$ | This work |
| pJL_sA20 | BeloBAC backbone, λ-red system, *cre*, *kan* and *rpoA* under strong promoter and $RBS_{20}$ | This work |
| pJL_sA40 | BeloBAC backbone, λ-red system, *cre*, *kan* and *rpoA* under strong promoter and $RBS_{40}$ | This work |
| pJL_sA60 | BeloBAC backbone, λ-red system, *cre*, *kan* and *rpoA* under strong promoter and $RBS_{60}$ | This work |
| pJL_sA80 | BeloBAC backbone, λ-red system, *cre*, *kan* and *rpoA* under strong promoter and $RBS_{80}$ | This work |
| pJL_sA98 | BeloBAC backbone, λ-red system, *cre*, *kan* and *rpoA* under strong promoter and $RBS_{98}$ | This work |

| | | |
|---|---|---|
| pJL_sAN | BeloBAC backbone, λ-red system, *cre*, *kan* and *rpoA* under strong promoter and $RBS_N$ | This work |
| pJL_nA20 | BeloBAC backbone, λ-red system, *cre*, *kan* and *rpoA* under native promoter and $RBS_{20}$ | This work |
| pJL_nA40 | BeloBAC backbone, λ-red system, *cre*, *kan* and *rpoA* under native promoter and $RBS_{40}$ | This work |
| pJL_nA60 | BeloBAC backbone, λ-red system, *cre*, *kan* and *rpoA* under native promoter and $RBS_{60}$ | This work |
| pJL_nA80 | BeloBAC backbone, λ-red system, *cre*, *kan* and *rpoA* under native promoter and $RBS_{80}$ | This work |
| pJL_nA98 | BeloBAC backbone, λ-red system, *cre*, *kan* and *rpoA* under native promoter and $RBS_{98}$ | This work |
| pJL_nAN | BeloBAC backbone, λ-red system, *cre*, *kan* and *rpoA* under native promoter and $RBS_N$ | This work |
| pJH05 | BeloBAC backbone, CEN/ARS, *his3*, *ura3*, λ-red system, *cre*, *kan* and RNAP operon with strong promoter, $RBS_{80}$ *rpoA* $RBS_N$ *rpoB* $RBS_{80}$ *rpoC* $RBS_{80}$ *rpoZ* | This work |

**Suppl. Table 4. List of primers**

| Primer name | Primer sequence | Description |
|---|---|---|
| RpoA fw | ATGCAGGGTTCTGTGACAGAG | Used for amplifying genes from *E. coli* genome |
| RpoA rv | TTACTCGTCAGCGATGCTTG | |
| RpoB fw | ATGGTTTACTCCTATACCGAG | |
| RpoB rv | TTACTCGTCTTCCAGTTCG | |
| rpoC fw | GTGAAAGATTTATTAAAGTTTCTG | |
| rpoC rv | TTACTCGTTATCAGAACCGCC | |
| rpoZ fw | ATGGCACGCGTAACTGTTCAG | |
| rpoZ rv | TTAACGACGACCTTCAGCAATAG | |
| rpoA+R20+spacer_fw | GACGTAATCGTCCAACTTTGAGTAGTGACACAATGCAGGGTTCTGTGACAGAG | Used for amplifying rpoA with respective RBSs |
| rpoA+R40+spacer_fw | GACGTAATCGTCCAACTTTGAGACATGACACAATGCAGGGTTCTGTGACAGAG | |
| rpoA+R60+spacer_fw | GACGTAATCGTCCAACTTTGCAAGAGGACACAATGCAGGGTTCTGTGACAGAG | |
| rpoA+R80+spacer_fw | GACGTAATCGTCCAACTTTGGGGAATGACACAATGCAGGGTTCTGTGACAGAG | |
| rpoA+R98+spacer_fw | GACGTAATCGTCCAACTTTGAGGAGTGACACAATGCAGGGTTCTGTGACAGAG | |
| rpoA+Rn+spacer_fw | GACGTAATCGTCCAACTTTGAGAGAGGACACAATGCAGGGTTCTGTGACAGAG | |
| Ps+spacer_fw | AGGCCGTGCCGGCACGTTGCAATACTTGACATATCACTGTGATTCACATATAATATGCGGACGTAATCGTCCAACTTTG | Used for amplifying RpoA with respective promoters |
| Pm+spacer_fw | AGGCCGTGCCGGCACGTTGCACCTATTGACAATTAAAGGCTAAAATGCTATAATTCCACGACGTAATCGTCCAACTTTG | |
| Pw+spacer_fw | AGGCCGTGCCGGCACGTTGCTCCCTTTGATATTGCATCCCGCGTATATAATATGTCGACGTAATCGTCCAACTTTG | |
| Pn+spacer_fw | AGGCCGTGCCGGCACGTTGCGATCGTCGAGCTTTACTCCAAGTAAAGCTTAGTACCAAAGAGACGTAATCGTCCAACTTTG | |
| rbs20+rpoB_fw | TGGGCATGCGCCTGGAAAACTGGCCACCGGCAAGCATCGCTGACGAGTAAGATGGCAACCCTATGGTTTACTCCTATACCGAG | Used for amplifying rpoB with respective RBSs |
| rbs40+rpoB_fw | TGGGCATGCGCCTGGAAAACTGGCCACCGGCAAGCATCGCTGACGAGTAAAGCTGGAACCCTATGGTTTACTCCTATACCGAG | |
| rbs60+rpoB_fw | TGGGCATGCGCCTGGAAAACTGGCCACCGGCAAGCATCGCTGACGAGTAACAAGAGAACCCTATGGTTTACTCCTATACCGAG | |
| rbs80+rpoB_fw | TGGGCATGCGCCTGGAAAACTGGCCACCGGCAAGCATCGCTGACGAGTAATAGAGAAACCCTATGGTTTACTCCTATACCGAG | |
| rbs98+rpoB_fw | TGGGCATGCGCCTGGAAAACTGGCCACCGGCAAGCATCGCTGACGAGTAATAGAGGAACCCTATGGTTTACTCCTATACCGAG | |
| rbsN+rpoB_fw | TGGGCATGCGCCTGGAAAACTGGCCACCGGCAAGCATCGCTGACGAGTAACTGAGGAACCCTATGGTTTACTCCTATACCGAG | |
| rpoB+overlap_rv | AAAAAAACCCCGCCGAAGCGGGGCGCCCAGTAGAAGCAGCAACTGTTAATTAATTACTCGTCTTCCAGTTCG | |
| rbs20+rpoC_fw | TGAAAGAGATTCGTTCGCTGGGTATCAACATCGAACTGGAAGACGAGTAAAGCTGCCAAATCCGTGAAAGATTTATTAAAGTTTCTG | Used for amplifying |

| | | |
|---|---|---|
| rbs40+rpoC_fw | TGAAAGAGATTCGTTCGCTGGGTATCAACATCGAACTGGAAGACGAGTAAAGCTGGCAAATCCGTGAAAGATTTATTAAAGTTTCTG | rpoC with respective RBSs |
| rbs60+rpoC_fw | TGAAAGAGATTCGTTCGCTGGGTATCAACATCGAACTGGAAGACGAGTAAAGTAGGCAAATCCGTGAAAGATTTATTAAAGTTTCTG | |
| rbs80+rpoC_fw | TGAAAGAGATTCGTTCGCTGGGTATCAACATCGAACTGGAAGACGAGTAAAAGGAGCAAATCCGTGAAAGATTTATTAAAGTTTCTG | |
| rbs99+rpoC_fw | TGAAAGAGATTCGTTCGCTGGGTATCAACATCGAACTGGAAGACGAGTAACGGAGGCAAATCCGTGAAAGATTTATTAAAGTTTCTG | |
| rbsN+rpoC_fw | TGAAAGAGATTCGTTCGCTGGGTATCAACATCGAACTGGAAGACGAGTAACGGGAGCAAATCCGTGAAAGATTTATTAAAGTTTCTG | |
| rpoC+overlap_rv | AAAAAAACCCCGCCGAAGCGGGGCGCCCAGTAGAAGCAGCAACTGTTAATTAATTACTCGTTATCAGAACCGCC | |
| rbs20+rpoz_fw | TTACTCGTTATCAGAACCGCCCAGACCTGCGTTCAGCAGTTCTGCCAGGCAGCTACTTTAAGTATGGCACGCGTAACTGTTCAG | Used for amplifying rpoZwith respective RBSs |
| rbs40+rpoZ_fw | TTACTCGTTATCAGAACCGCCCAGACCTGCGTTCAGCAGTTCTGCCAGGCAGACATTTTAAGTATGGCACGCGTAACTGTTCAG | |
| rbs60+rpoZ_fw | TTACTCGTTATCAGAACCGCCCAGACCTGCGTTCAGCAGTTCTGCCAGGCACGAGTTTTAAGTATGGCACGCGTAACTGTTCAG | |
| rbs80+rpoZ_fw | TTACTCGTTATCAGAACCGCCCAGACCTGCGTTCAGCAGTTCTGCCAGGCAGGCACTTTAAGTATGGCACGCGTAACTGTTCAG | |
| rbs98+rpoZ_fw | TTACTCGTTATCAGAACCGCCCAGACCTGCGTTCAGCAGTTCTGCCAGGCAGGAGCTTTAAGTATGGCACGCGTAACTGTTCAG | |
| rbsN+rpoZ_fw | TTACTCGTTATCAGAACCGCCCAGACCTGCGTTCAGCAGTTCTGCCAGGCGAGCTTTTTAAGTATGGCACGCGTAACTGTTCAG | |
| rpoZ+T+HR_rv | ACTCCGTTACAAAGCGAGGCTGGGTATTTCCCGGCCTTTCTGTTATCCGCAAAAAAACCCCGCCGAAGCGGGGCGCCCAGTAGAAGCAGCAACTGTTAATTAATTAACGACGACCTTCAGCAATAG | RpoZ reverse with terminator |
| pHLUM_OH fw | CCTTTTACAGCCAGTAGTGCTCGCCGCAGTCGAGCGACAGGGCGAAGCCCGATCGCTTGCCTGTAACTTACACGC | Used for amplifying *his3* and CEN/ARS |
| pHLUM_OH rv | GTGTGTAAGCAGAATATATAAGTGCTGTTCCCTGGTGCTTCCTCGCTCACTAATCGGTGTCACTACATAAGAAC | |
| Bac_fw | TACCGCTGAAAGTTCTGCAAAG | pBelobac11 backbone amplification |
| Bac_rv | AACGTGCCGGCACGG | |
| pSC020_OH_fw | TTACCCAGGCCGTGCCGGCACGTTGCGGCCGCGGATAACAGAAAGGCCGG | Amplification of λ-red and *cre* |
| pSC020_OH_rv | TGAGTTTTTCTAATTGCGTTGCGCTCACTG | |
| Ura3_OH_fw | CATATTCCATTTTGTAATTTCGTGTCGTTTCTATTATGAATTTCATTTATCACTCAACCCTATCTCGGTC | Amplification of *ura3* |
| Ura3_OH_rv | ACTCCGTTACAAAGCGAGGCTGGGTATTTCCCGGCCTTTCTGTTATCCGCTCAATTCATCATTTTTTTTTTATTCTTTTTTTTG | |
| rpoA_KO_fw | GATCGTCGAGCTTTACTCCAAGTAAAGCTTAGTACCAAAGAGAGGACACATAGAGTCGGACTTCGCGTTCGC | Amplification of KO template for *rpoA* |
| rpoA_KO_rv | GATGGCGCATGACCTTATCCTTCTCAGTAAAACCTTAACCTGTGATCCGGGCTAGTTATTGCTCAGCGGTGG | |
| rpoB_KO_fw | GTCCGCTCAATGGACAGATGGGTCGACTTGTCAGCGAGCTGAGGAACCCTTAGAGTCGGACTTCGCGTTCGC | Amplification of KO |

| | | |
|---|---|---|
| rpoC_KO_rv | CATAAAAAAACCCGCCGAAGCGGGTTTTTACGTTATTTGCGGATTAACGAGCTAGTTATTGCTCAGCGGTGG | template for *rpoB* and *rpoC* |
| rpoZ_KO_fw | GATTTCAGTATCATGCCCAGTCATTTCTTCACCTGTGGAGCTTTTTAAGTTAGAGTCGGACTTCGCGTTCGC | Amplification of KO template for *rpoZ* |
| rpoZ_KO_rv | ATCAGTTGATTCAGGCTTTCAAACAGATACAAGGGCGACCCGCTTTGTGAGCTAGTTATTGCTCAGCGGTGG | |
